# Diroximel fumarate acts through Nrf2 to attenuate methylglyoxal-induced nociception in mice and decreases ISR activation in DRG neurons

**DOI:** 10.1101/2023.12.22.572877

**Authors:** Muhammad Saad Yousuf, Marisol Mancilla Moreno, Jiahe Li, Lucy He, Danielle Royer, Jennifer Zhang, Brodie J Woodall, Peter M Grace, Theodore J Price

## Abstract

Diabetic neuropathic pain is associated with elevated plasma levels of methylglyoxal (MGO). MGO is a metabolite of glycolysis that causes mechanical hypersensitivity in mice by inducing the integrated stress response (ISR), which is characterized by phosphorylation of eukaryotic initiation factor 2α (p-eIF2α). Nuclear factor erythroid 2-related factor 2 (Nrf2) is a transcription factor that regulates the expression of antioxidant proteins that neutralize MGO. We hypothesized that activating Nrf2 using diroximel fumarate (DRF) would alleviate MGO-induced pain hypersensitivity. We pretreated male and female C57BL/6 mice daily with oral DRF prior to intraplantar injection of MGO (20 ng). DRF (100 mg/kg) treated animals were protected from developing MGO-induced mechanical and cold hypersensitivity. Using *Nrf2* knockout mice we demonstrate that Nrf2 is necessary for the anti-nociceptive effects of DRF. In cultured mouse and human dorsal root ganglion (DRG) sensory neurons, we found that MGO induced elevated levels of p-eIF2α. Co-treatment of MGO (1 µM) with monomethyl fumarate (MMF, 10, 20, 50 µM), the active metabolite of DRF, reduced p-eIF2α levels and prevented aberrant neurite outgrowth in human DRG neurons. Our data show that targeting the Nrf2 antioxidant system with DRF is a strategy to potentially alleviate pain associated with elevated MGO levels.

**Perspective:** This study demonstrates that activating Nrf2 with DRF prevents the development of pain caused by MGO in mice and reduces ISR in mouse and human DRG *in vitro* models. We propose that Nrf2 activators like DRF should be tested to alleviate diabetic neuropathic pain associated with elevated MGO in patients.

**Graphical Abstract:** 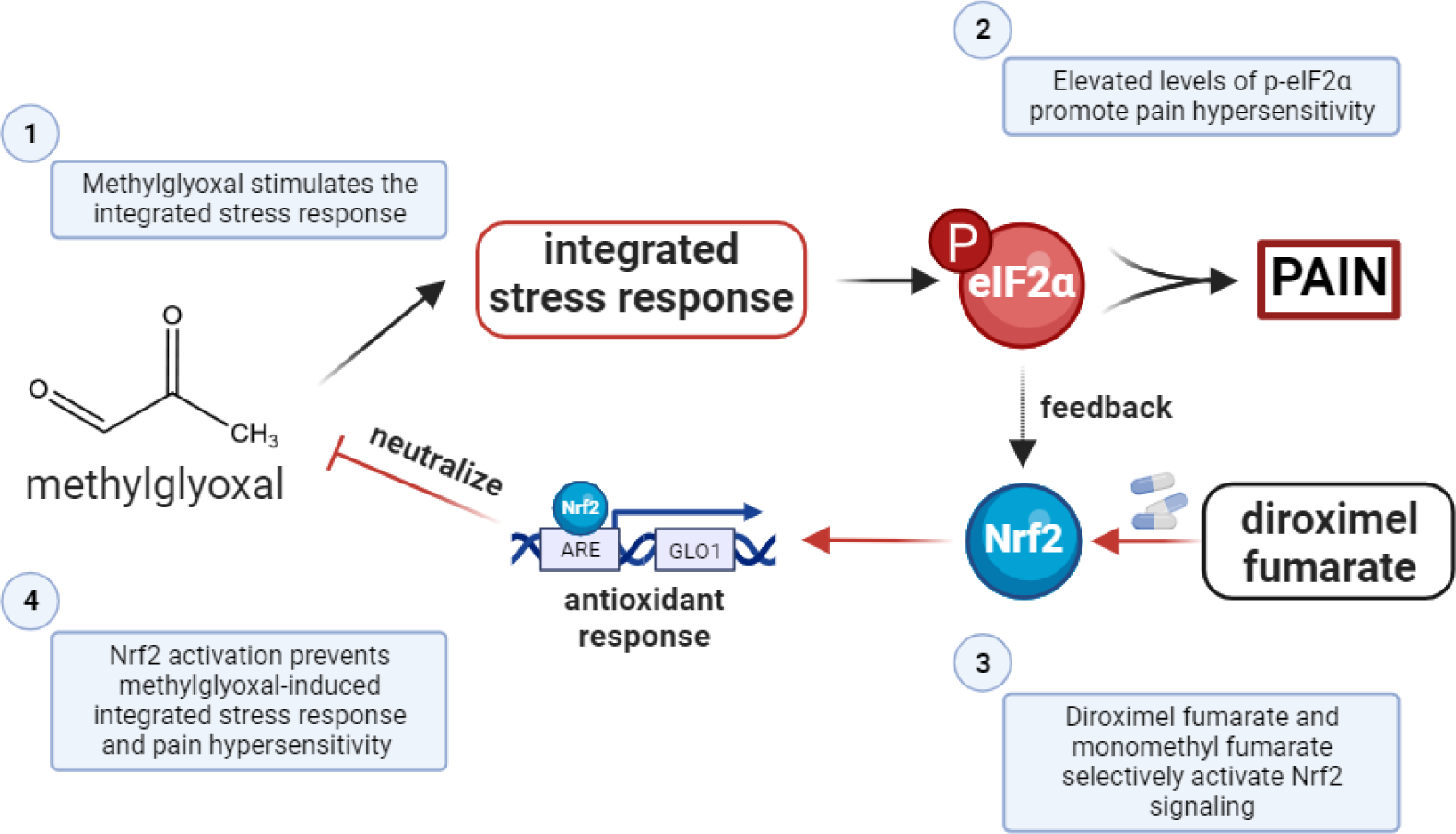

**Article Highlights:**

- MGO induces mechanical and cold hypersensitivity in mice that is prevented with pre-treatment with DRF.
- DRF pre-treatment does not protect Nrf2-knockout mice from developing pain hypersensitivity suggesting that Nrf2 is necessary for DRF’s antinociceptive effects.
- MMF, the active metabolite of DRF, prevents MGO-induced increase in p-eIF2a levels in mouse and human DRG neurons *in vitro*.
- MMF prevents MGO-induced aberrant neurite outgrowth in human DRG neurons.
- Nrf2 activators, like the FDA-approved DRF, is an option to alleviate neuropathic pain in patients with diabetes.

## Introduction

Diabetes is the most common metabolic disorder in the world, affecting around ten percent of the population (1). Recent estimates have found that approximately half of all diabetic individuals eventually develop neuropathy making diabetic neuropathic pain the most common form of neuropathic pain (2). Diabetic neuropathic pain is associated with worsening quality of life and increased burden on healthcare (2). Currently diabetic neuropathic pain is treated using anticonvulsants like gabapentin and pregabalin, antidepressants like duloxetine, and opioids like tapentadol and tramadol (2; 3). However, these therapies have modest efficacy and serious side-effects (3–6).

Prior studies have shown that oxidative stress and antioxidant pathways play a crucial role in the development and maintenance of diabetic neuropathy (7; 8). Methylglyoxal (MGO) is a reactive dicarbonyl metabolite of glycolysis associated with elevated blood glucose levels, inflammation, and aging (9). Individuals suffering from diabetic neuropathic pain have elevated plasma levels of MGO (1 µM), which is approximately three times higher than levels in diabetics without neuropathic pain (10; 11). MGO interacts with amino acid residues like cysteine, arginine, and lysine to produce advanced glycation end-products (AGEs) and promote the synthesis of reactive oxygen species (12–14). We have recently demonstrated that levels of MGO (1 µM) associated with diabetic neuropathic pain stimulates the integrated stress response (ISR) and that pharmacologically inhibiting the ISR alleviates MGO-induced mechanical hypersensitivity (15; 16). The ISR is an adaptive response to stressors, like accumulated misfolded proteins and elevated reactive oxygen species, that is initiated by the phosphorylation of eukaryotic initiation factor 2α (eIF2α) (17–19). The ISR suppresses global protein synthesis and facilitates the translation of specific mRNAs, like the mRNA encoding activating transcription factor 4 (ATF4), which influence neuronal activity and viability (19). The ISR also activates nuclear factor erythroid 2-related factor 2 (Nrf2), a potent transcription factor, to protect cells from and reverse oxidative stress (20; 21). However, ISR-mediated Nrf2 antioxidative response is not sufficient to prevent pain hypersensitivity.

The glutathione system acts as a critical node in the antioxidant response because dicarbonyls like MGO are first neutralized by a non-enzymatic reaction with glutathione to generate hemithioacetal which is further metabolized by glyoxalase 1 (Glo1) and glyoxalase 2 (Glo2) to generate inert D-lactate (22). Nrf2 is a transcription factor that regulates the expression of many antioxidant proteins by binding to their antioxidant response elements (ARE) (23). The genes for glutamate cysteine ligase (GCL), the rate limiting step in the formation of glutathione, and Glo1 contain an ARE in their 5’ untranslated regions and their transcription is promoted by Nrf2 (23; 24). Fumaric acid esters like diroximel fumarate (DRF), dimethyl fumarate (DMF), and their active metabolite, monomethyl fumarate (MMF), are known activators of the Nrf2 antioxidant pathway (25–27). All are FDA-approved for treatment of relapsing remitting multiple sclerosis (27; 28). Thus, we hypothesized that DRF and the Nrf2 antioxidant pathway can be leveraged to reduce MGO-induced nociceptive hypersensitivity and signaling associated with pain in both mouse and human DRG neurons.

Inhibiting the production of reactive oxygen species or promoting the antioxidant response have both been postulated to treat patients with diabetic neuropathy (7; 29; 30). However, a safe and effective therapeutic has remained elusive. Our results provide support for targeting the Nrf2 signaling pathway as a strategy for alleviating pain associated with MGO-associated neuropathy.

## Research Design and Methods

### Animals

Wild-type C57BL/6 mice were obtained from Charles River and a colony was maintained at UT Dallas. Nrf2KO mice (B6.129X1-Nfe2l2^tm1Ywk^/J) were obtained from Jackson Laboratories (Strain # 017009) and cross-bred with wild-type C57BL/6 mice over multiple generations. A colony of Nrf2KO and wild-type littermates was maintained at UT Dallas where behavior experiments were performed. Genotyping was performed by Transnetyx (Memphis, Tennessee) based on protocol and PCR primers outlined by Jackson Laboratories (Protocol # 26266). Animals were maintained on a 12-hour light-dark cycle. Adult male and female mice at least 12 weeks old were used throughout the study. All experiments were performed according to guidelines established by the National Institutes of Health and the Animal Care and Use Committee at the University of Texas at Dallas (protocol # 14-04).

### Mouse samples – DRG cultures

Lumbar (L1-L6) mouse DRGs were extracted from wild-type male (n=2) and female (n=2) mice. DRGs from male and female mice were processed and cultured together. DRGs were digested in pre-warmed (37°C) 3 ml enzyme mix (2 mg/mL STEMzyme I (Worthington Biochemical #LS004107), 4 µg/mL DNAse I (Worthington Biochemical #LS002139) in Hanks’ Balanced Salt Solution (HBSS) without calcium and magnesium (Gibco #14170161)). Enzyme mix and DRGs were incubated at 37°C in a shaking water bath. DRG tissue was triturated 2-3x every 20 minutes until tissue was completely homogenous. Cells were passed through 70 µm cell strainer and then layered onto 1 ml of 15% bovine serum albumin (BSA BioPharm #71-040-002) in HBSS. This gradient was centrifuged at 900g for 5 min at room temperature. The pellet was resuspended in prewarmed media (DMEM/F12+GlutaMax (Gibco #10565-018), 1% Pen-Strep (ThermoFisher #15070063) and 1% N2-A (STEMCell #07152)). DRG neurons were plated on poly-D-lysine (70,000-150,000 Da) (Sigma-Aldrich #P6407) coated coverslips.

### Human samples – DRG cultures

Human DRGs were recovered from organ donors in collaboration with Southwest Transplant Alliance. Human DRGs (UNOS ID AJHC468, 29-year-old male) were harvested and cultured according to previously outlined protocols (16; 31). In brief, human DRGs were microdissected into 1-2mm chunks. Human DRG neurons were cultured on 100 ug/ml poly-D-lysine (>300,000 Da) coated 12 mm coverslips and allowed to equilibrate in vitro for 24 hours prior to being treated with MGO and MMF for the next 24 hours. Cells were fixed with 10% formalin and processed for immunocytochemistry.

### Drugs and Chemicals

Methylglyoxal (∼40%) was obtained from Sigma-Aldrich (M0252). It was further dissolved in 0.9% normal saline or culturing media for *in vivo* and *in vitro* experiments, respectively. Diroximel fumarate (DRF) (Biogen) was dissolved in 10 mM citric acid / 0.5% carboxymethylcellulose / 0.02% Tween80 adjusted to pH 5.0. For *in vitro* studies, monomethyl fumarate (MMF) (Sigma-Aldrich # 651419) was initially dissolved to 500 mM in 100% DMSO. Subsequent dilutions of MMF were performed in culturing media.

### Mouse behavior testing

Animals were accustomed to human touch four days prior to any testing. Baseline behavior testing was performed for two (2) consecutive days prior to any drug treatment. Animals were habituated on mesh racks for one (1) hour before being tested. Experimenters were blind to genotype and treatment conditions of the animals. Only the left paw was injected with vehicle or MGO and only that paw was tested throughout the experiment.

Mechanical paw withdrawal thresholds were obtained based on the Simplified Up-Down protocol of von Frey filaments test (32). Von Frey filaments (Ugo Basile) were calibrated using a weigh scale (VWR, 314AC) to 0.01 grams. Cold sensitivity was assessed using acetone applied to the plantar surface of the injected paw using a transfer pipette (Fisher, 13-711-20). Nocifensive behaviors (i.e. vigorous shaking and licking) were timed for up to 45 seconds using a stopwatch. The injured paw was tested three times, and the response times were averaged.

### Immunocytochemistry (ICC)

Immunocytochemistry protocol was based on a previously published report (16). In brief, cultured mouse and human DRG neurons were fixed with 10% formalin (Thermo-Fisher, 23-245684) at room temperature for 10 minutes followed by 3 washes of 1X PBS. Cells were subsequently blocked for 1 hour at room temperature with 10% normal goat serum (NGS, R&D systems S13150H) dissolved in 1X PBS-Tx (0.02% Triton X-100 in 1X PBS, Sigma-Aldrich X100-5ML). Cells were incubated overnight at 4°C in primary antibody dissolved in antibody solution (2% NGS and 2% bovine serum albumin (BSA) dissolved in PBS-Tx). Cells were washed three times with 1X PBS. Secondary antibodies were also dissolved in antibody solution. Cells were incubated in secondary antibodies for 1 hour at room temperature followed by three 1X PBS washes. DAPI (Cayman Chemicals, 14285) was dissolved in 1X PBS and washed twice with 1X PBS. Human DRGs were incubated with TrueBlack (Biotium 23007) for 30 seconds to quench lipofuscin signal. Coverslips were mounted onto slides before being imaged. Confocal images were obtained on Olympus FV-1200 microscope at 10X magnification. Antibodies used in this study were: p-eIF2α (1:500, Cell Signaling #3398), β3 tubulin (1:1000, Sigma #T8578), DAPI (1:10,000, Cayman Chemicals #14285), Alexa Fluor 488 (1:500, Life Technologies, A11034), and Alexa Fluor 555 (1:500, Life Technologies, A21428).

### Sholl Analysis

Sholl analysis was performed using a previously established protocol using the Neuroanatomy plugin according to author recommendations (16; 33). Only solitary cells were used for analysis. β3-tubulin staining was used to mark neurons and neurites. Cells were imaged at 20X magnification on a Olympus FV3000 confocal and Z-stacked at 1µm intervals. The area under the curve was calculated using GraphPad Prism 10.

### Data Analysis

All results are presented as mean ± standard error of mean (SEM). Statistical analyses are mentioned in figure legends. In general, comparisons between two groups were performed using Student’s t-test with or without Welch’s correction depending on the homoscedasticity of the dataset. When comparing more than two conditions, one-way or two-way ANOVA’s were used with or without correction for homoscedasticity of the dataset. BioRender was used to build schematics. Data was graphed and analyzed on GraphPad Prism 10.

## Results

### DRF prevents MGO-induced mechanical and cold hypersensitivity

In this study, we hypothesized that DRF would prevent MGO-induced mechanical and cold pain hypersensitivity. DRF activates Nrf2 and the glyoxalase and glutathione antioxidant pathways which are key to neutralizing MGO (**Fig 1A**). We treated male and female mice daily with vehicle (DRF-vehicle) or DRF at either 60 mg/kg (DRF-60) or 100 mg/kg (DRF-100) given orally for 5-days. On day 3, an intraplantar injection of MGO (20 ng) was also administered one hour after oral DRF for that day. von Frey assay and acetone tests were performed to assess mechanical and cold sensitivity (**Fig 1B**). After the first two days of DRF treatment, we found no change in mechanical thresholds in male and female mice in response to only DRF administration at either 60 mg/kg and 100 mg/kg, suggesting that DRF has no intrinsic effects on mechanical thresholds. Male and female mice that were administered MGO and treated with DRF-vehicle developed hypersensitivity to tactile stimulation when compared to vehicle-only animals (**Fig 1C, D**). DRF treatment, particularly at 100 mg/kg (DRF-100), prevented MGO-induced mechanical pain hypersensitivity. Area under the curve of von Frey thresholds were used to directly compare each treatment group (**Fig 1E, F**). Female animals were protected at both the 60 mg/kg and 100 mg/kg doses while the male animals were only benefited from a higher dose of 100 mg/kg of the drug. DRF treatment at both 60 mg/kg and 100 mg/kg completely prevented cold hypersensitivity on day 3 in both male and female mice (**Fig 1G, H**). By day 5, cold sensitivity returned to baseline levels in all conditions. These results show that treatment with DRF prevents mechanical and cold hypersensitivity in response to MGO.

**Figure 1.**
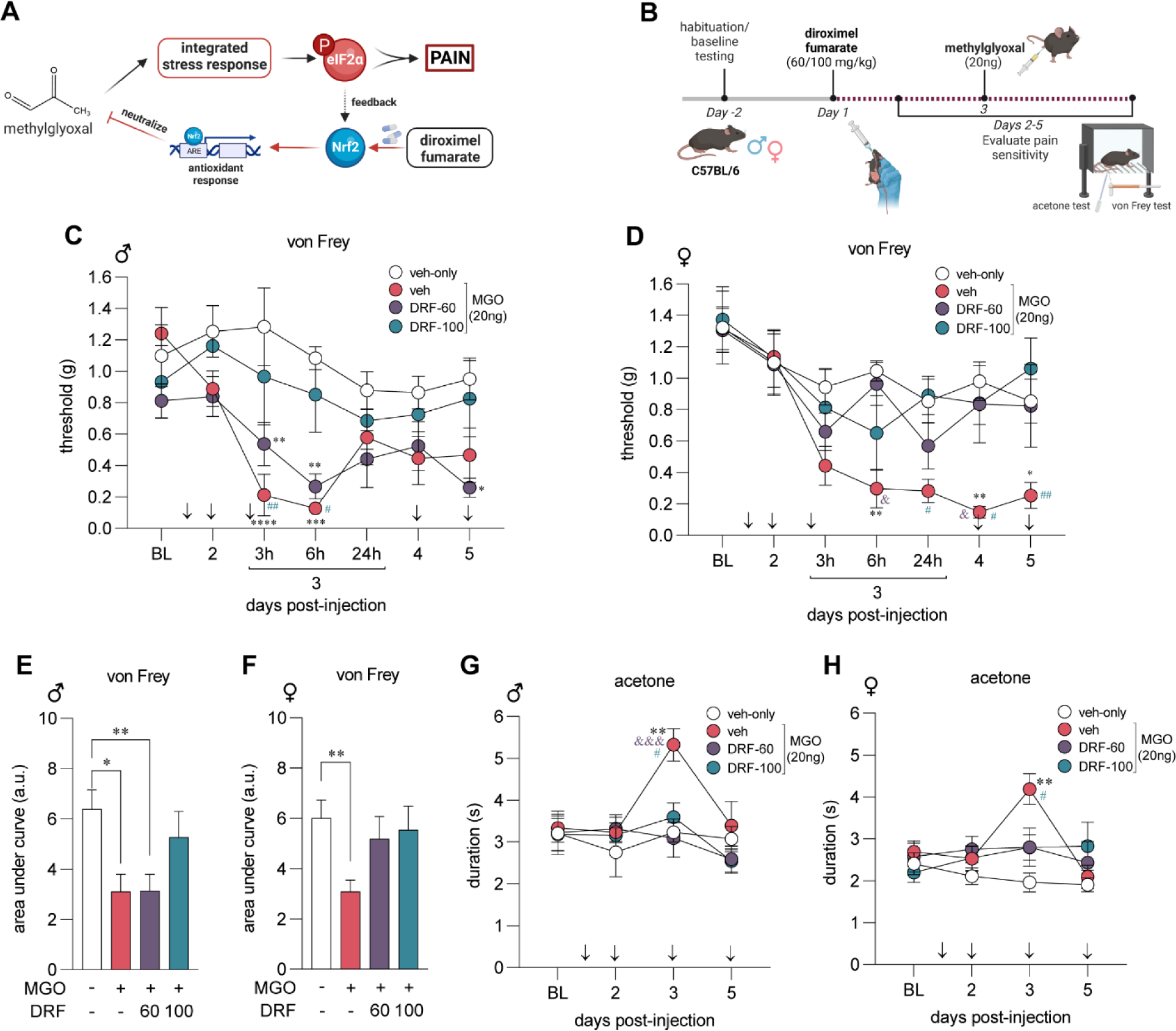
Diroximel fumarate (DRF) pretreatment protects against methylglyoxal (MGO)-induced pain hypersensitvity. (A) MGO stimulates the integrated stress response (ISR) and promotes the phosphorylation of eIF2α. DRF is a prodrug and is metabolized to MMF which activates the Nrf2 antioxidant response neutralizing MGO. (B) Male (n=8 per condition) and female (n=8 per condition) mice were treated with DRF (60 or 100 mg/kg) for five days. MGO (20 ng) was administered in an intraplantar injection on the third day, 1-hour after oral DRF administration for the day. Mechanical and cold sensitivity was assessed using von Frey and acetone tests, respectively. Acetone test was performed an hour after von Frey testing was complete. We observed that male (C) and female (D) mice had reduced von Frey thresholds in response to MGO injection on day 3. Daily treatment with DRF prevented MGO-induced mechanical hypersensitivity following MGO injection on day 3. (E, F) Using area under the curve of von Frey thresholds, we determined that DRF at a dose of 100 mg/kg were effective at preventing MGO-induced tactile hypersensitivity in males while both 60 mg/kg and 100 mg/kg doses were effective in females. MGO produced cold hypersensitivity in male (G) and female (H) mice which was abrogated in DRF treated mice at both doses. For (C, D, G, H) significance was calculated with repeated measures two-way ANOVA with Tukey’s multiple comparison test. *, &, # p<0.05; **, ## p<0.01; ***, &&& p<0.001; **** p<0.0001. * indicates comparison to veh-only, & indicates comparison to MGO+DRF-60, # indicates comparison to MGO+DRF-100. For (E, F) signficance was caluclated with a one-way ANOVA with Brown-Forsythe correction. * p<0.05; ** p<0.01.

### DRF requires Nrf2 to counteract the pain promoting effects of MGO

Endoplasmic reticulum (ER) pathology is a canonical feature of elevated MGO (15; 34). MGO stimulates PERK to phosphorylate eIF2α and Nrf2 to instigate the ISR and antioxidant mechanisms, respectively. We have previously shown that p-eIF2α signaling is critical for developing mechanical hypersensitivity in multiple models of neuropathic pain, including MGO-induced hypersensitivity (15). To delineate whether the protective effects of Nrf2 arm affects the duration and intensity of nociceptive hypersensitivity following MGO administration, we treated wild-type and global Nrf2KO animals with a single intraplantar injection of MGO (20ng) or vehicle (saline) and followed these animals for 20 days. Since we found no evidence of sex-specific effects of DRF in **Fig 1**, we performed these experiments with an equal number of male (n=4) and female (n=4) mice in each group. We found that MGO induced robust tactile hypersensitivity in wild-type (n=8) and Nrf2KO (n=8) animals as compared to the vehicle treated group and that effect was comparable between genetic strains (**Fig 2A, B**). We directly compared the response to MGO using area under the curve of von Frey thresholds of each genotype and found no significant difference in the response of Nrf2KO and wild-type mice to MGO administration (**Fig 2C**).

**Figure 2.**
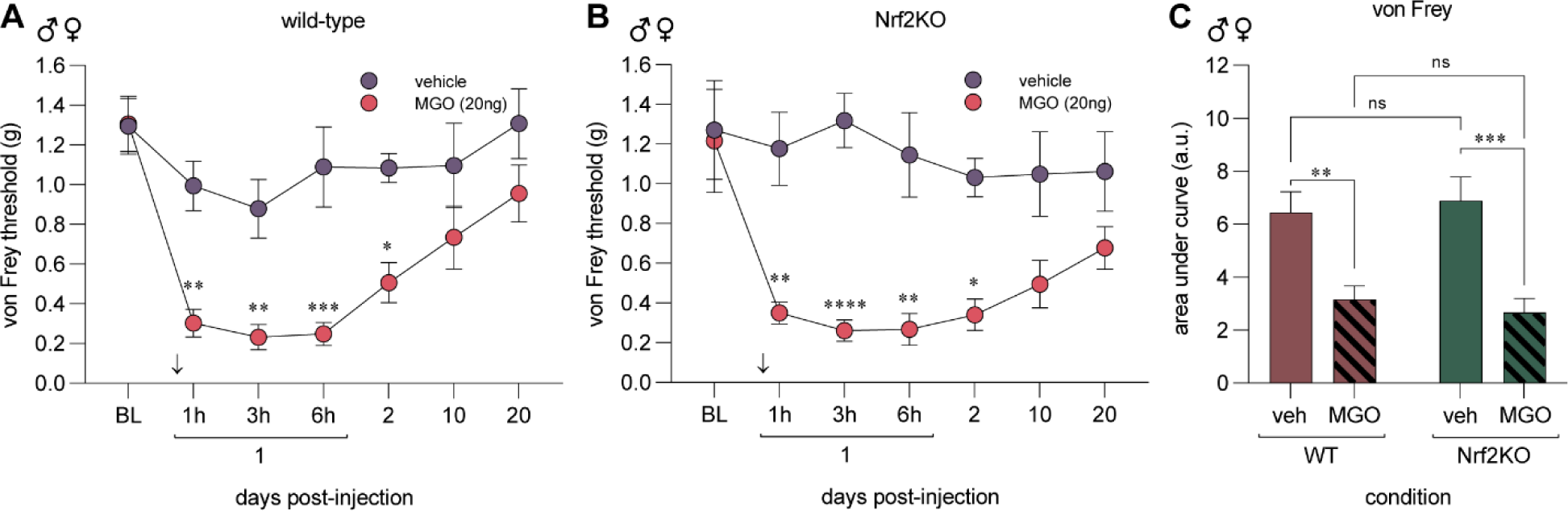
MGO induces mechanical pain hypersensitivity in Nrf2KO mice. Wild-type (A) and Nrf2KO (B) animals were injected MGO (20ng) or vehicle (saline) in the left paw. Mechanical sensitivity was assessed using the von Frey test for up to 20 days after injection. MGO (20ng) injection produced mechanical hypersensitivity in both wild-type and Nrf2KO mice. (C) Area under the curve of von Frey thresholds was used to directly compare the response of wild-type and Nrf2KO animal to MGO injection. Each condition included 4 male and 4 female mice for a total of 8 animals. For (A, B) statistical analysis was performed with repeated measures two-way ANOVA followed by Sidak multiple comparison test. * p<0.05, ** p<0.01, ***p<0.001, ****p<0.0001. In (C), a two-way ANOVA with Tukey’s test was used. ** p<0.01, ***p<0.001.

We next sought to answer whether DRF’s anti-nociceptive effects are mediated specifically by Nrf2. We treated male and female Nrf2KO mice (n=8) and their wild-type littermates (n=8) with 100 mg/kg of DRF (DRF-100) and MGO following the same treatment paradigm as outlined in **Fig 1**. We discovered that Nrf2KO animals developed MGO-induced mechanical hypersensitivity that was not prevented with DRF treatment (**Fig 3**). On the contrary, wild-type littermates were protected from MGO-induced tactile hypersensitivity (**Fig 3**). Our data taken from **Fig 2** and **Fig 3** suggests that Nrf2 is not necessary for the development, maintenance, or resolution of MGO-induced mechanical hypersensitivity while pharmacologically promoting Nrf2 signaling plays an important role in preventing it.

**Figure 3.**
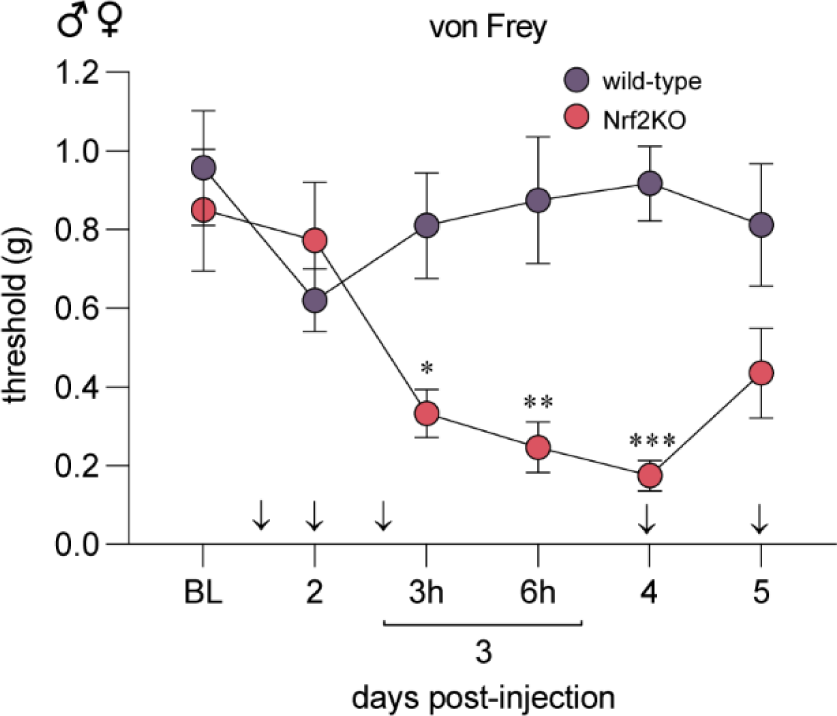
DRF treatment fails to prevent mechanical hypersensitivity in Nrf2KO mice. Wild-type and Nrf2KO animals were treated with DRF at 100 mg/kg based on the paradigm outlined in **Fig 1A**. MGO (20 ng) was injected into the paw on day 3 since the start of oral DRF treatment in wild-type (n=8, 4 males and 4 females) and Nrf2KO (n=8, 4 males and 4 females) littermates. Nrf2KO animals developed mechanical hypersensitivity compared to the wild-type animals suggesting that DRF treatment was unable to prevent MGO-induced hypersensitivity in Nrf2KO animals. A repeated measures two-way ANOVA was used followed by Sidak’s post hoc test. *p<0.05, **p<0.01, ***p<0.001.

### MMF suppresses MGO-induction of the ISR and reduces p-eIF2α in mouse and human sensory neurons

DRF is a prodrug that is metabolized in the small intestines to the active compound, MMF, and an inactive compound, 2-hydroxyethyl succinimide (HES) (27; 35). We have previously shown that MGO promotes the ISR by phosphorylating eIF2α and that reversing or preventing the ISR, alleviates pain associated with MGO (15; 16). To investigate whether MMF inhibits the ISR, we treated sensory neuron cultures from independent male (n=2) and female (n=2) mouse dorsal root ganglia (DRG) with MGO (1 µM) and MMF (10, 20, 50 µM) or vehicle for 24 hours. Afterwards cells were fixed with formalin and processed for immunohistochemistry. We observed an increase in p-eIF2α immunoreactivity in cells treated with MGO as compared to vehicle-treated cells (**Fig 4A, B**). This increase in p-eIF2α levels was prevented when MGO treated cultures were co-treated with MMF in a concentration-dependent manner (**Fig 4C, D**).

**Figure 4.**
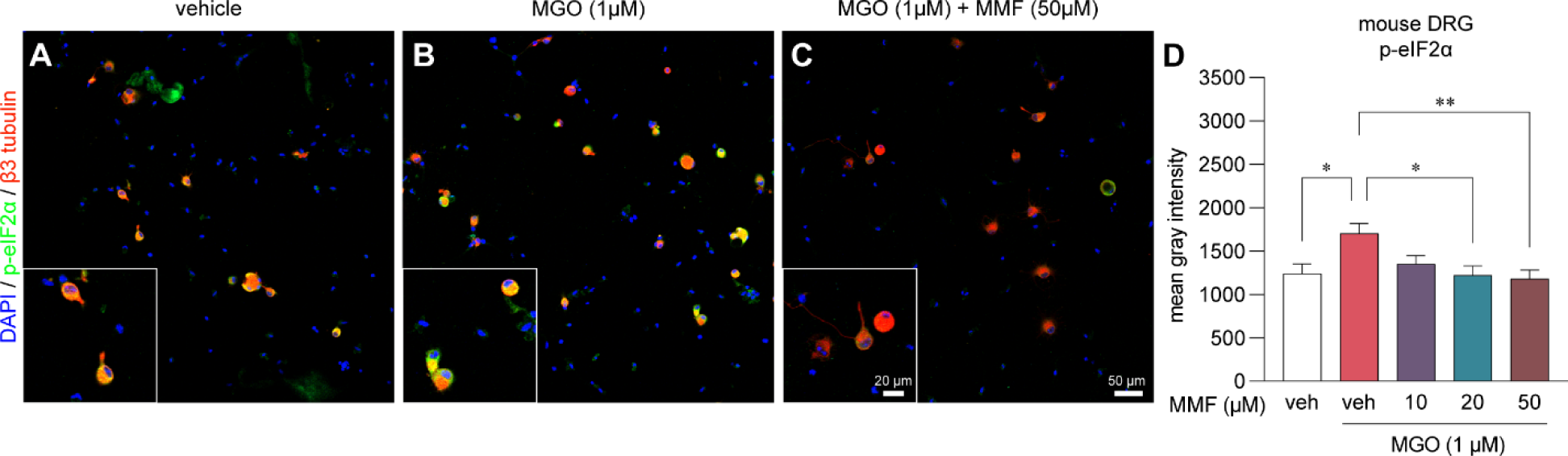
Monomethyl fumarate (MMF) prevents MGO-evoked increases in p-eIF2α levels in mouse DRG neurons. (A-C) Representative images of immunohistochemistry of mouse DRG neurons cultured for 24 hours and subsequently treated with vehicle-only, MGO (1µM), or MGO (1µM) plus MMF (10, 20, 50 µM). (D) We observed a significant increase in the levels of p-eIF2α when cells were treated with MGO as compared to vehicle-treated cells. MMF treatment with MGO prevented elevation of p-eIF2a particularly at 20 µM and 50 µM concentrations. Neurons were identified by their expression of beta3 tubulin. Vehicle-only (n=18), MGO (n=37), MGO+10µM MMF (n=20), MGO+20µM MMF (n=16), MGO+50µM MMF (n=32). A one-way ANOVA was used to calculate significance. *p<0.05, **p<0.01.

To expand the clinical relevance of our findings, we sought to replicate our murine findings in human sensory neurons from cultured DRGs recovered from organ donors (**Fig 5A-C**). MGO (1 µM) treatment elevated p-eIF2α immunoreactivity in human sensory neurons and cotreating them with MGO (1 µM) and MMF (10, 20, 50 µM) prevented the increase in p-eIF2α levels (**Fig 5D**). Similar to mouse DRG neurons, we observed a significant reduction in p-eIF2α immunoreactivity at 20 and 50 µM MMF concentrations (**Fig 5D**). Hence, mouse and human DRG neurons *in vitro* are protected from MGO-induced ISR by MMF.

**Figure 5.**
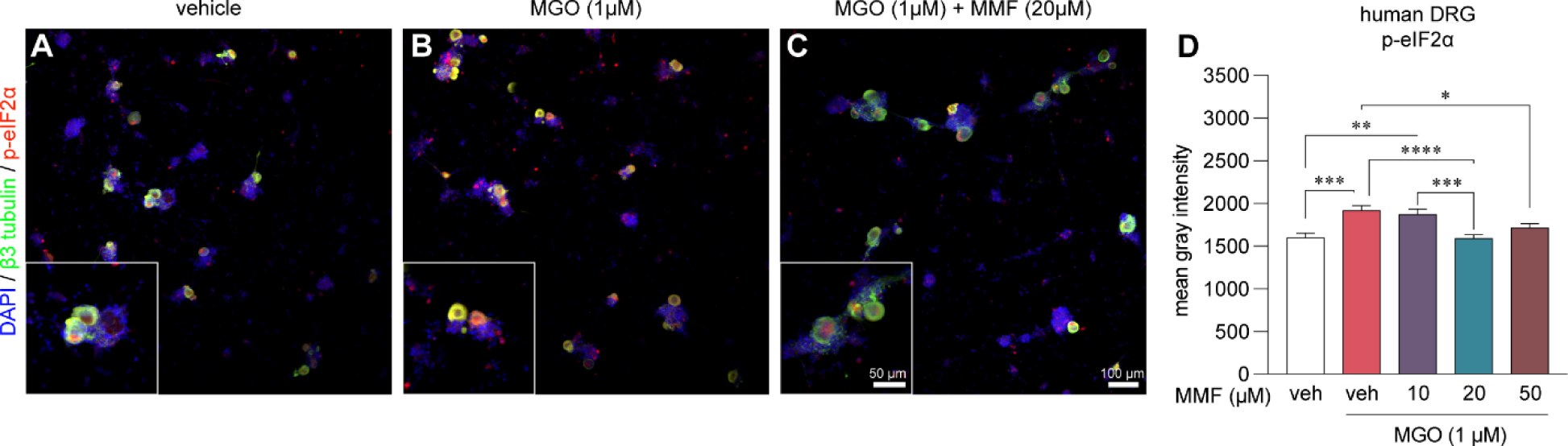
Monomethyl fumarate (MMF) prevents MGO-evoked increases in p-eIF2α levels in human DRG neurons. (A-C) Representative images of human DRG neurons obtained from organ donors treated with vehicle, MGO (1µM), and MGO plus MMF. (D) MGO (1µM) increases p-eIF2α in human DRG neurons which is prevented with 20 µM and 50 µM MMF cotreatment. Vehicle-only (n=170), MGO (n=152), MGO+10µM MMF (n=125), MGO+20µM MMF (n=176), MGO+50µM MMF (n=111). A one-way ANOVA was used to calculate statistical significance. *p<0.05, **p<0.01, ***p<0.001, ****p<0.0001.

### MMF reverses aberrant neurite outgrowth in human DRG neurons

Changes in axon innervation targets, axonal integrity, and axonal regeneration potential are directly associated with diabetic neuropathy (36; 37). We measured neurite outgrowth by Sholl analysis as a functional readout of MGO-induced effects on DRG axons and we tested for potential protective effects of MMF in human DRG neurons. We treated human DRG neurons with vehicle, MGO (1µM), or MGO (1µM) with MMF (10, 20, 50 µM) for 24 hours. Cells were subsequently fixed and immunolabeled for β3 tubulin as a marker for neurons and their neurites (**Fig 6A-C**). The number of intersections of all neurites of each cell were quantified and compared. We found that MGO treatment of human DRGs for 24 hours stimulated neurite outgrowth with increased neuronal complexity and arborization as compared to vehicle-only treated cells (**Fig 6D, E**). We further discovered that co-treatment with MMF prevented aberrant arborization caused by MGO in a concentration dependent manner (**Fig 6D, E**). Sholl analysis of MGO and MMF-treated cells remained identical to cells treated with vehicle alone. We conclude that MGO influences molecular ISR signaling and neurite outgrowth of human DRG neurons *in vitro* and these effects can be prevented by activating Nrf2 signaling using MMF.

**Figure 6.**
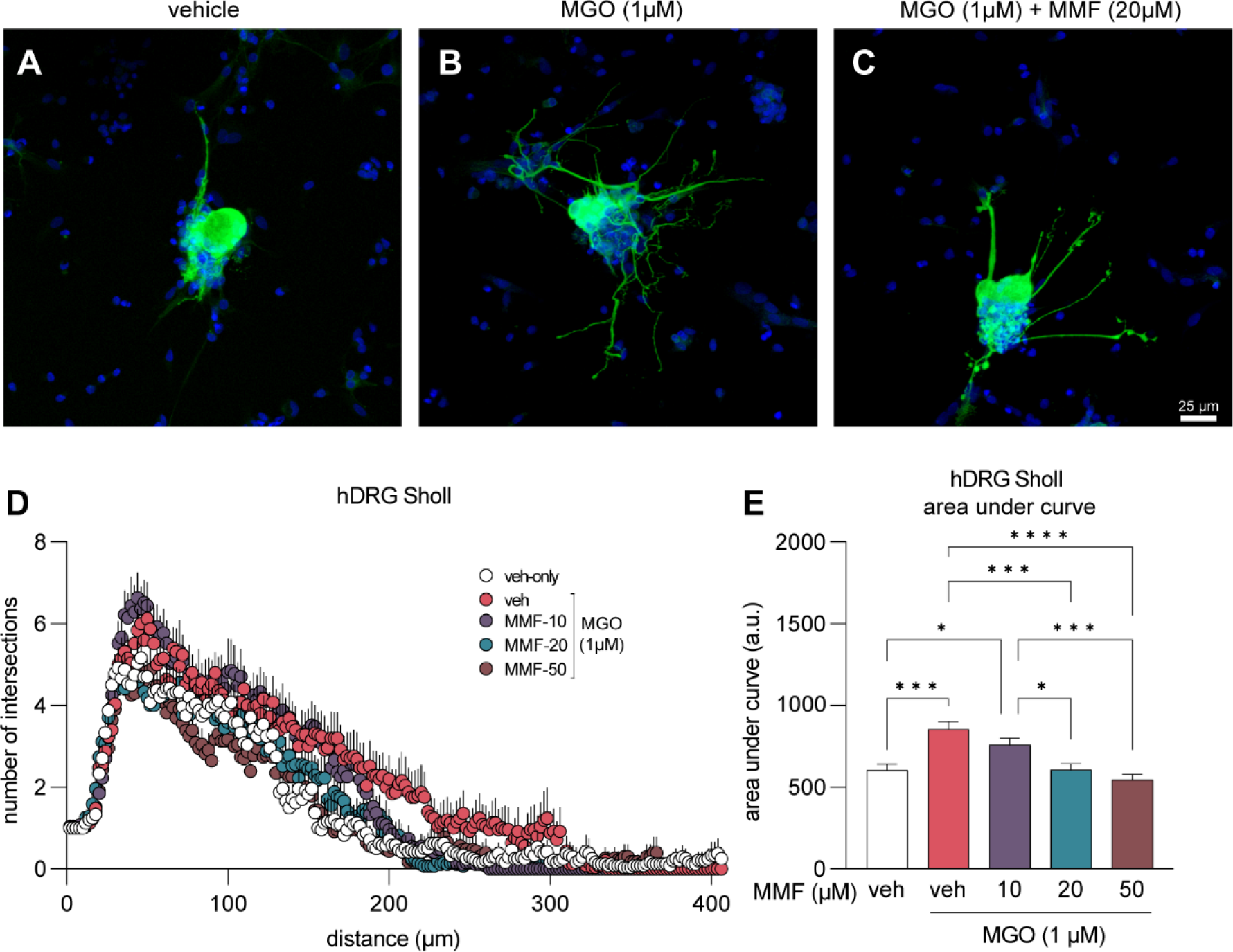
Neurite outgrowth induced by MGO is prevented with MMF treatment. (A-C) Example images of human DRG neurons immunolabeled with β3 tubulin (green) to visualize neuronal cell body and nerites. Neurons were treated with vehicle, MGO (1µM), or MGO (1µM) plus MMF (10, 20, 50 µM) for 24 hours prior to fixation and immunostaining. (D) Sholl analysis was performed to quantify neurite outgrowth and complexity. (E) Area under the curve of Sholl analysis shows that MMF treatment prevents MGO-induced neurite outgrowth, particularly at 20 µM and 50 µM concentrations. Vehicle-only (n=24), MGO (n=25), MGO+10µM MMF (n=27), MGO+20µM MMF (n=27), MGO+50µM MMF (n=23). A one-way ANOVA was used to calculate statistical significance. *p<0.05, ***p<0.001, ****p<0.0001.

## Discussion

In this study, we demonstrate that DRF prevents MGO-induced mechanical and cold hypersensitivity in mice. Using Nrf2KO mice, we demonstrate that endogenous Nrf2 signaling in response to MGO (1 µM) does not affect the initiation, maintenance, or resolution of MGO-induced mechanical pain hypersensitivity. However, the antinociceptive effect of DRF is lost in Nrf2KO mice suggesting that DRF’s effects on Nrf2 are essential for reducing mechanical hypersensitivity in mice. At a concentration of MGO observed in the plasma of people with diabetic neuropathic pain (1 µM), MGO activates the ISR and elevates levels of p-eIF2α (15; 16). This effect is significantly blunted in cultured mouse and human DRG neurons when MGO is co-administered with the active metabolite of DRF, MMF. Furthermore, we show that MGO treatment at 1 µM stimulates aberrant neurite outgrowth that is prevented with MMF treatment in a concentration-dependent manner in human DRG neurons. Our data collectively provides evidence for targeting the Nrf2 antioxidant system for treating MGO-induced nociception and suggests that drugs targeting this pathway, like DRF, could be amenable for pain treatment in people with diabetic neuropathy and elevated MGO levels.

Oxidative stress is a causative factor in the progression of diabetic neuropathy (7; 8; 38; 39). Here, we demonstrate that the Nrf2 signaling pathway can be leveraged to fortify and protect against ongoing oxidative and dicarbonyl stress responsible for MGO-induced pain hypersensitivity. Nrf2 is a master transcription factor that regulates the transcription of other antioxidant genes, particularly those with an ARE element in their promoter region (23). Nrf2 activators, DMF and monoethylfumarate, in cultured human astrocytes engage the glutathione and glyoxalase pathways that peak 12 to 24 hours after administration (40). We thus hypothesized that pretreatment with DRF will prime the antioxidant system such that exogenous MGO is promptly neutralized, protecting the cells against the detrimental effects of MGO. Although not directly addressed in our work, this hypothesis is testable in future experiments.

The interaction between the ISR and the Nrf2 pathway is complex with feedforward and feedback mechanisms. Conditions of ER stress prime PERK to phosphorylate both eIF2α and Nrf2, dissociating Nrf2 from its inhibitor Keap1, and facilitating its nuclear translocation (21). A recent report also found that ER stress-induced ATF4 increases *Nrf2* transcription, increasing the global pool of Nrf2 to counteract reactive oxygen species (20). In addition, MGO at high concentrations (1 mM) can covalently modify Keap1 to form dimers and promote Nrf2 activity (41). Based on these prior observations, we expected that MGO-induced mechanical hypersensitivity would be more pronounced or take longer to resolve in Nrf2KO mice than WT animals. To determine how endogenous Nrf2 signaling affects the characteristics of MGO-induced mechanical hypersensitivity, we treated Nrf2KO mice with MGO and assessed their mechanical hypersensitivity for up to 20 days. Surprisingly, we found no difference in tactile hypersensitivity between wild-type animals and Nrf2KO animals. This suggests that the endogenous activation of Nrf2 antioxidant systems is not sufficient to prevent or alleviate pain hypersensitivity and that an exogenous application of Nrf2 activators is required to galvanize the antioxidant pathway. One possible reason for this is that the kinetics of the ISR and Nrf2 antioxidant response are different. ER stress and ROS potently activate the ISR in the first few hours while Nrf2-mediated antioxidant response requires roughly 24 hours (20). Coupled with the observation that exogenous MGO induces pain hypersensitivity in the first few hours, it is likely that ISR-dependent pain-causing mechanisms have already been set in motion before the endogenous antioxidant Nrf2 response takes effect.

Fumaric acid esters obtained from fungi have been used to treat inflammatory conditions since medieval times (27). Next generation fumaric acid esters were developed in the late 1980s (27). Most recently, DRF received FDA approval for relapsing remitting MS (28). In clinical trials DRF was found to be superior to its predecessor, dimethyl fumarate, due to its improved side effect profile, particularly a reduction in adverse gastrointestinal effects and abdominal pain because of a lack of methanol by-production created by esterase activity (35). We have previously shown that DMF’s anti-nociceptive effects are dependent on Nrf2 activity in rodent model of traumatic nerve injury (26). Since DMF and DRF share the same active metabolite, MMF, we speculated that DRF’s anti-nociceptive effects are specifically due to its activity on Nrf2. A recently published study found that DRF changed the transcriptome of cultured human astrocytes that was consistent with Nrf2 activation however no causal link was presented (25). Our data provide evidence that DRF’s antinociceptive effects in the MGO-induced pain model are specifically mediated by Nrf2 as DRF was unable to prevent MGO-induced mechanical hypersensitivity in global Nrf2KO mice. Contrary to our findings, DMF reduced pain hypersensitivity in a rodent model of complex regional pain syndrome through an Nrf2-independent mechanism (42). This suggests that the specificity of DMF/DRF-Nrf2 signaling may depend on the pain model being studied.

While our data provides promising results for an Nrf2-activator in reducing the pain-promoting effects of MGO, it comes with some limitations. First, we acknowledge that our pain behavior data is not coupled with molecular analysis of harvested tissue from treated animals. We have yet to demonstrate that DRF treatment reduces ISR and p-eIF2α levels *in vivo* and that this coincides with the antinociceptive effect of DRF, however, our in vitro experiments in mice and human neurons provide clear support for this hypothesis. Second, our experiments with DRF were prevention experiments where we pretreated animals with DRF prior to the MGO injection. In a clinical setting, preventative pain medicines are rarely prescribed. Whether DRF can reverse already established MGO-induced pain hypersensitivity remains to be tested. Because DRF is clinically available, we contend that the in vivo and in vitro data we provide is strong support for planning of a future clinical trial.

Since MGO can be measured in patient populations, our work provides a strong rationale for testing Nrf2 activators in people with diabetic neuropathic pain specifically associated with elevated MGO concentrations. This approach is feasible because levels of MGO (43) and MGO-specific AGEs like MG-H1 and CEL can easily be assessed from routine blood chemistry analysis of diabetic patients, stratifying a patient population that would ideally respond to DRF treatment. Having been FDA-approved, DRF has already demonstrated itself to be safe and well-tolerated so such clinical trials could be conducted.

## Acknowledgments

The authors are grateful to the organ donors and their families for the gift of life and research provided by their organ donation. The authors thank Anna Cervantes, Geoffrey Funk, and Peter Horton of the Southwest Transplant Alliance for human tissue recovery.

## Author Contributions

MSY designed and performed experiments and analyzed data. MMM, LH, JZ, DR, BJW performed experiments. JL generated Nrf2-knockout animals and maintained the colony. PMG and TJP supervised the study. MSY is the guarantor of this work and takes responsibility for the contents of this article.

